# Abuse liability, antinociceptive, and discriminative stimulus properties of IBNtxA

**DOI:** 10.1101/2020.05.30.125450

**Authors:** Ariful Islam, Mohammad Atiqur Rahman, Megan B. Brenner, Allamar Moore, Alyssa Kellmyer, Harley Buechler, Frank DiGiorgio, Vincent Verchio, Laura McCracken, Mousumi Sumi, Robert Hartley, Joseph R. Lizza, Gustavo Moura-Letts, Bradford D. Fischer, Thomas M. Keck

## Abstract

**Rationale:** IBNtxA (3-iodobenzoyl naltrexamine) is a novel μ opioid receptor (MOR) agonist structurally related to the classical MOR antagonist naltrexone. Recent studies suggest IBNtxA preferentially signals through truncated MOR splice variants, producing a unique pharmacological profile resulting in antinociception with reduced side effects, including no conditioned place preference (CPP) when tested at a single dose. IBNtxA represents an intriguing lead compound for preclinical drug development targeting truncated MOR splice variants but further evaluation of its in vivo pharmacological profile is necessary to evaluate its potential.

**Objective:** The purpose of this study was to independently verify the antinociceptive properties of IBNtxA and to more completely examine the rewarding properties and discriminative stimulus effects of IBNtxA. These results will allow broader assessment of IBNtxA as a translational candidate or lead compound for further development.

**Results:** IBNtxA was synthesized and compared to morphine in a variety of mouse behavioral assays. 3 mg/kg IBNtxA was equipotent to 10 mg/kg morphine in a hot plate analgesia assay. In drug discrimination testing using mice trained to discriminate between 3 mg/kg IBNtxA and DMSO/saline vehicle, the κ agonist U-50488 fully substituted for IBNtxA. Classical μ agonist morphine, δ agonist SNC162, NOP agonist SCH 221510, and μ/NOP partial agonist buprenorphine each partially substituted for IBNtxA. IBNtxA up to 3 mg/kg did not produce a place preference in CPP. Pretreatment with 3 mg/kg IBNtxA but not 1 mg/kg IBNtxA attenuated acquisition of place preference for 10 mg/kg morphine. 3 mg/kg IBNtxA attenuated morphine-induced hyperlocomotion but did not alter naloxone-precipitated morphine withdrawal.

**Conclusions:** Overall IBNtxA has a complicated opioid receptor pharmacology *in vivo*. These results indicate that IBNtxA produces potent antinociception and has low abuse liability, likely driven by substantial κ agonist signaling effects.

## Introduction

Opioid drugs are critical medications used to treat acute and chronic pain (Ballantyne and Mao 2003; McQuay et al. 1997; Sullivan and Howe 2013). The safety and tolerability of opioid analgesics is severely limited due to side effects that include sedation, constipation, respiratory depression, and abuse liability (Ahlbeck 2011; Ling et al. 1984; Ling et al. 1985; Ling et al. 1983; Pasternak 2014; Pasternak and Pan 2013). The dramatic recent growth in prescription opioid misuse and abuse has contributed to a major public health emergency, often referred to as the opioid epidemic (Coussens et al. 2019; Manchikanti et al. 2012; Volkow et al. 2014). In 2016, nearly 62 million people in the United States either filled or refilled opioid prescriptions at least once (Centers for Disease Control and Prevention 2017), and an estimated 11.8 million people misused or abused prescription opioids and/or the illicit opioid heroin (SAMSHA 2017). These data indicate a pressing need for new analgesics with reduced abuse liability.

Several strategies have been pursued in the search for analgesic medications with reduced abuse liability, including drugs that target not only the μ-opioid receptor (MOR), but also the δ-opioid receptor (DOR), κ-opioid receptor (KOR), and/or nociceptin/orphanin opioid receptor (NOR) (Kiguchi et al. 2020; Spahn and Stein 2017; Turnaturi et al. 2019). Partial agonists, such as buprenorphine, provide analgesic effects with reduced abuse liability in part due to their low efficacy (Gudin and Fudin 2020; Walsh et al. 1995). Additionally, formulations that include antagonists (e.g., naloxone and naltrexone) have been designed to reduce the abuse potential of opioid drugs (Mendelson and Jones 2003; Setnik et al. 2019).

A promising class of molecules, β-naltrexamine analogues of naltrexone, were synthesized in 2011 and determined to have somewhat surprising agonist activity despite the inclusion of the cyclopropylmethyl moiety found on naltrexone and other classical opioid receptor antagonists (Majumdar et al. 2011a). One compound in this series, 3-iodobenzoyl naltrexamine (IBNtxA), was later pharmacologically characterized and reported to have an analgesic potency 10-fold greater than that of morphine, while producing reduced side effects typical of classic MOR agonist analgesics, including no observable respiratory depression or withdrawal symptoms, limited inhibition of gastrointestinal transit, and no rewarding behavior in a single-dose conditioned place preference (CPP) assay (Majumdar et al. 2011b). A series of follow-up studies tested the hypothesis that IBNtxA preferentially signals via selected MOR splice variants, particularly exon 11-associated variants that result in a truncated MOR protein with only six transmembrane regions (6TM/E11), which results in its unique pharmacological profile (Grinnell et al. 2016; Grinnell et al. 2014; Lu et al. 2015; Majumdar et al. 2011b; Majumdar et al. 2012; Marrone et al. 2016; Wieskopf et al. 2014; Xu et al. 2009). These studies provided evidence that IBNtxA-mediated analgesia could be primarily a product of 6TM/E11 MOR activation, and additionally determined that 6TM/E11 MOR variants contribute to the analgesic effects for certain MOR agonists, such as fentanyl, buprenorphine and heroin, but not for morphine (Grinnell et al. 2016; Lu et al. 2015; Majumdar et al. 2011b; Pan et al. 2009). It is not currently known to what degree signaling by 6TM/E11 splice variants mediates drug-seeking and -taking behaviors for opioid drugs or contributes to the discriminative stimulus properties of opioid ligands.

As the functional roles of 6TM/E11 MOR splice variants are not well understood, pharmacological tools like IBNtxA can be used to investigate the biological effects of 6TM/E11 MOR signaling. The purpose of this study was to independently verify the analgesic properties of IBNtxA and to examine more completely the rewarding properties and discriminative stimulus effects of IBNtxA; these studies will allow us to better understand the *in vivo* effects of IBNtxA, more thoroughly evaluate its antinociceptive properties, abuse liability, and provide a necessary foundation for future studies to directly investigate the consequences of 6TM/E11 MOR signaling.

## 2. Materials and Methods

### Animals

In total, the following experiments used 209 adult male C57BL/6 mice obtained from Charles River Laboratories (Wilmington, MA). All experiments were performed within the vivarium at Cooper Medical School of Rowan University (CMSRU). Mice were group-housed (four animals per cage) in polycarbonate cages in a temperature- and humidity-controlled vivarium with *ad libitum* access to food and water with enrichment provided by paper Bio-Huts and/or nestlets. Housing maintained a standard light/dark cycle (lights on at 07:00, lights off at 19:00), therefore all experiments were performed during the light part of the cycle in separate procedure rooms from the housing facility. Animals were weighed daily and monitored for general health and behavioral parameters, including potential signs of significant opioid withdrawal. All experiments were conducted in accordance with the Guide for the Care and Use of Laboratory Animals (US National Academy of Sciences) and were approved by the Institutional Animal Care and Use Committee of Rowan University. The CMSRU animal facility is provisionally accredited by the Association for Assessment and Accreditation of Laboratory Animal Care International.

### Drugs

IBNtxA was synthesized at Rowan University using the procedures described below. Morphine sulfate was purchased from Henry Schein (Melville, NY). Cocaine HCl was purchased from Sigma Aldrich (St. Louis, MO). Buprenorphine, naltrexone, U-50488, SCH 221510, and SNC 162 were purchased from Tocris (Minneapolis, MN).

All drugs were delivered via intraperitoneal (i.p.) injections at a volume of 10 mL/kg. Drug dilutions were premixed to provide a given mg/kg dose when given an injection volume scaled to mouse body weight, measured prior to every test. Cocaine HCl was dissolved in physiological saline. All other drugs used a common vehicle of 10% dimethyl sulfoxide (DMSO) and 90% physiological saline, henceforth referred to as DMSO vehicle. To solubilize all drugs, they were initially completely dissolved in 100% DMSO and diluted to their final concentration with sterile physiological saline.

### IBNtxA synthesis

A novel synthesis of IBNtxA was developed using naltrexone as starting material. Detailed methods are available in the supplementary material. Briefly, IBNtxA was synthesized using commercial naltrexone (Tocris) converted into naltrexamine, and reacted with 2,5-Dioxopyrrolidin-1-yl 3-iodobenzoate. Purified IBNtxA HCl was then used for all experiments.

### Behavioral Procedures

#### Hot plate analgesia test

##### Justification

The hot plate test is a widely used test to evaluate the antinociceptive potential of analgesic medications. The test relies upon the natural nocifensive responses of rodents when placed on a hot metallic plate, and the latency to those responses induced by analgesics, including opioids and nonsteroidal anti-inflammatory drugs (NSAIDs) (Bannon and Malmberg 2007; Deuis et al. 2017; Le Bars et al. 2001; Woolfe and MacDonald 1944). Because analgesic effects in the hot plate assay are mediated supraspinally, this experiment allowed for identification and characterization of analgesic, and centrally active, doses of IBNtxA for further testing.

##### Apparatus

The hot plate analgesia test was performed using methods derived from previously published reports (Bannon and Malmberg 2007; Deuis et al. 2017). The hot plate analgesia meter for small laboratory animals (Columbus Instruments) was used for this test featured an aluminum surface controlled via a built-in digital thermometer to maintain surface temperatures to within 0.1 °C and a digital timer with 0.1 sec precision. The square surface plate was enclosed by a clear acrylic cage to confine animals during testing. Morphine is well-characterized in the hot plate assay and served as our positive control. Before and after all training or testing procedures, the apparatus was cleaned with 70% isopropyl alcohol and allowed to dry.

##### Treatment groups

Twenty-four drug-naïve male C57BL/6 mice were used for this study, separated into three groups of *n* = 8: 10 mg/kg morphine, 1 mg/kg IBNtxA, and 3 mg/kg IBNtxA.

##### Testing

The hot plate was set to continuously provide a 55 °C temperature during testing. Depression of the start/stop button for the digital timer signaled the beginning and end of each experiment and the elapsed time for each test was recorded manually. Animals were placed on the heated surface and two independent observers looked for behavioral changes to indicate painful sensations, including licking or fluttering of paws or jumping. Upon witness of these behaviors, the timer was stopped and animals were removed immediately from the chamber. To prevent injury, all experiments had a maximum hot plate exposure time of 20 seconds and animals were immediately removed at this time if they had not yet shown a pain response. The latency time after placing mice on the metallic hot plate provided the threshold level of animals.

Prior to drug administration, each mouse was weighed and tested twice, 15 minutes apart, to establish baseline nocifensive response times. Immediately following the second baseline test, drugs were administrated via i.p. injection. The first post-injection test occurred 15 minutes after injection and animals were re-tested every 15 minutes for 60 minutes or until the latency time returned to baseline. Data normalization was performed by taking the two pre-injection latency times and averaging them for each animal as a baseline (BL). A % Max Possible Effect was calculated using the formula: 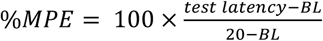.

#### Drug Discrimination

##### Justification

Drug discrimination (DD) allows the study of neuropharmacological and behavioral effects of psychoactive drugs, using the effects of a delivered drug as a discriminative stimulus paired with reinforcers (e.g., food). Animals are trained to press one of two levers to obtain food after receiving drug injections, and to press the other lever to obtain food after vehicle injections. After sufficient training, subjects can be used to determine whether a test substance is “like” or “unlike” the drug used for training (Colpaert 1999; Solinas et al. 2006; Young 2009), allowing evaluation of the subjective effects of IBNtxA in comparison to other opioid ligands. These procedures were adapted from Solinas et al. (2006).

##### Apparatus

DD training and testing used mouse operant chambers (Med Associates, Fairfax, VT) placed in sound attenuating cabinets and connected to a computer running MED-PC software (version 4). Each operant chamber featured two nose poke holes equipped with infrared beams. A liquid dipper was located between the two nose poke holes, connected via tubing to a pump and syringe that discharged vanilla Ensure for 3 seconds (delivering an approximate 0.1 mL volume) as a palatable food reward when animals earned a programmed reward. All rewards were accompanied by light and tone stimuli during the duration of the 3-second reward delivery.

##### Treatment groups

Eight drug-naïve male C57BL/6 mice were used for this study and received all initial drug discrimination training. One mouse did not meet inclusion criteria and was excluded from further testing. One mouse died near the end of the experiment, therefore two doses of SCH 221510 (0.3 and 1.0 mg/kg) and two doses of cocaine (3 and 10 mg/kg) were tested on only six trained animals.

##### Food restriction

Mice had *ad lib* access to water, but were food restricted for approximately 22 hours before each training or testing session. Mice were weighed daily and were kept within 10% of their free-feeding weight. Following all training or testing sessions, animals were given *ad lib* access to chow for 2 hours.

##### Training

Mice were trained initially to nose poke for Ensure rewards using a fixed-ratio 1 (FR1) schedule, in which a single nose poke on either side initiated reward delivery and associated cues. Following successful nose poke training, in which mice received at least 90 of 100 possible rewards in a 1-hour time period, animals were trained to discriminate between DMSO vehicle (10% DMSO and 90% saline) and 3 mg/kg IBNtxA. During the training phase, mice received i.p. injections of 3 mg/kg IBNtxA or DMSO vehicle and were placed in the operant chamber 15 minutes prior to the start of training, with the start of the session indicated by a house light turning on. IBNtxA and vehicle were given with a pseudorandom order of training to avoid day-of-the-week training effects. For all chambers, IBNtxA was programmed to be associated with the right nose poke hole, and DMSO vehicle with the left nose poke hole. In order to earn an Ensure reward, animals were required to complete an unbroken FR response in the correct nose poke hole. Over the course of training, the FR requirement was increased until mice were correctly nose poking >90% on a FR5 schedule; mice were considered to have successfully learned the DD procedure at a given FR level when ≥ 90% of the initial 10 nose pokes in a given training session matched the desired response. All training sessions lasted for 60 minutes or until 100 rewards were earned. Before and after all training and testing procedures, each test apparatus and floor insert was cleaned with 70% isopropyl alcohol and allowed to dry completely.

##### Testing

After meeting FR5 training criteria, mice were tested with varying doses of IBNtxA (0.33-3.0 mg/kg), morphine (0.33-10 mg/kg), U-50488 (0.33-10 mg/kg), buprenorphine (0.10-1.0 mg/kg), SNC162 (3-18 mg/kg), SCH 221510 (1-10 mg/kg) and cocaine (3-10 mg/kg) given via i.p. injection. For each test, mice were given a drug injection and placed in the operant chamber 15 minutes prior to the start of testing session, with the start of the session indicated by a house light turning on. Drug discrimination was measured by the first response (drug side or vehicle side) after the start of the session, after which the session was immediately ended with no rewards given. This limited, stringent testing procedure was adopted after initial studies determined that IBNtxA discrimination training was easily disrupted by rewards earned while exposed to some tested drugs (possibly owing to IBNtxA being a relatively weak discriminative stimulus). This procedure allowed much quicker (∼1 week) re-establishment of drug discrimination between tests, but it also results in data that are less typical of DD reports in the literature: instead of reporting the proportion of overall nose pokes (drug-paired vs. vehicle-paired) following testing, we report the proportion of animals whose intial responses were on the drug-paired vs. vehicle-paired side. Likewise, because we cannot report standard drug effects on response rates (e.g., nose pokes/sec), we report time to initial nose poke at the start of the session.

#### Conditioned Place Preference (CPP)

##### Justification

The basic principle underlying CPP—widely used to evaluate the subjective effects of drugs of abuse (Bardo and Bevins 2000; Bardo et al. 1995a; Tzschentke 2007)—is that primary reinforcers (e.g., legal or illicit drugs, food, water, or sex) are paired with contextual stimuli, which acquire secondary reinforcing properties via Pavlovian contingency (Agustin-Pavon et al. 2008; Bardo et al. 1995b; Liu et al. 2008). Procedures for CPP experiments were adapted from previously published protocols (Carboni and Vacca 2003; Cunningham et al. 2006; Prus et al. 2009; Roux et al. 2001). Morphine is well-characterized in rodent CPP models (Hall et al. 2003; Karami and Zarrindast 2008; Koek 2016; Shippenberg et al. 1996; Vindenes et al. 2008) and served as our positive control.

##### Apparatus

CPP studies were performed using modular open field apparatuses separated into three compartment via black Plexiglas™ place preference chamber inserts (Stoelting, Wood Dale, IL). These chambers featured two 18 × 20 cm rectangular test compartments connected via a 10 × 20 cm central compartment. The two test compartments were differentiated by wall markings (vertical green stripes vs. white circles) and by custom-made, 3D-printed plastic floor inserts (yellow with circular holes, paired with stripe side, vs. blue with a shallow square grid, paired with circle side) printed at Rowan University. Test compartments could be closed off during training using a removable black Plexiglas™ blocker placed in the central compartment.

Four test chambers were arrayed on a vertical shelving unit, two per shelf. Over each chamber was a small fluorescent light (overall illumination ∼700 lux) and a USB camera connected to a computer running Any-maze software (version 4.99). Between every animal, and before and after all training and testing procedures, each test apparatus and floor insert was cleaned with 70% isopropyl alcohol and allowed to dry completely.

##### Treatment groups

120 male drug-naïve C57BL/6 mice were used for this study. Following initial preference training, animals were divided into seven groups, each receiving a different potential reinforcer dose for the drug-paired side and DMSO vehicle for the vehicle-paired side: DMSO vehicle (*n* = 17), 3 mg/kg morphine (*n* = 14), 10 mg/kg morphine (*n* = 22), 20 mg/kg morphine (*n* = 12), 0.3 mg/kg IBNtxA (*n* = 13), 1 mg/kg IBNtxA (*n* = 16), and 3 mg/kg IBNtxA (*n* = 14). Preliminary studies with a higher 5.6 mg/kg dose of IBNtxA were abandoned due to extreme behavioral disruptions, including loss of motor coordination.

##### Initial preference

Before preference training, animals were tested for an initial side preference (i.e., a preference for either test compartment) by placing naïve mice in the chamber with full access to all compartments for 30 minutes. Initial compartment preference was calculated as the ratio of time spent in one test compartment over the total time spent in both test compartments; time spent in the central compartment was recorded but not included in this calculation. Twelve animals with >65% initial preference for one test compartment were excluded from further study. Consistent with an unbiased experimental procedure (Tzschentke 2007), the drug-paired and vehicle-paired compartments for each animal were assigned regardless of a given animal’s initial preference score using a Latin square design, which evenly distributed compartment pairing, box location, and 1^st^-day treatment across all treatment groups. Vehicle-only animals were assigned the striped side as their “drug” compartment and the circle side as their “vehicle” compartment.

##### Drug conditioning

Conditioning (*a.k.a.* training or acquisition) occurred during ten training sessions over ten days. During conditioning, animals received drug or vehicle injections immediately prior to being placed and confined in the designated test compartment for 30 minutes. Drug and vehicle exposures alternated daily.

We chose an unbiased CPP procedure because while biased approaches can provide greater sensitivity to detect drug effects that depend on baseline motivational states of the animal, or to detect anxiolytic and anti-aversive drug effects, they also have a higher false-positive rate, which we wished to avoid (Cunningham et al. 2003; Cunningham et al. 2006; Tzschentke 1998; 2007).

##### Preference test

Trained mice were tested for side preference in a CPP expression trial in which subjects received no injection and were free to access entire apparatus (i.e., both drug- and vehicle-paired contexts) for 30 minutes.

##### Analysis

A preference score was calculated for the both the initial preference test (Δ_initial_) and for the final preference test (Δ_final_) by subtracting the seconds spent in the vehicle-paired compartment was from the seconds spent in the drug-paired compartment (time in the central compartment was ignored). An overall preference score of a given drug dosage was calculated by using the formula ΔΔsec = Δ_final_ -Δ_initial_.

#### Locomotor activity

##### Justification

To determine whether IBNtxA might induce substantial locomotor effects, and/or alter morphine-induced locomotor behavior, we tested whether 3 mg/kg IBNtxA would alter locomotor activity in an open field.

##### Apparatus

Locomotor studies were performed using four 40 × 40 cm modular open field apparatuses (Stoelting, Wood Dale, IL) arrayed on a vertical shelving unit, two per shelf. Over each chamber was a small fluorescent light (overall illumination ∼700 lux) and a USB camera connected to a computer running Any-maze software (version 4.99). Between every animal, and before and after all training and testing procedures, each test apparatus and floor insert was cleaned with 70% isopropyl alcohol and allowed to dry completely.

##### Treatment groups

41 male drug-naïve C57BL/6 mice were used for this study. Animals were divided into four groups, each receiving a pre-treatment and a post-treatment: DMSO vehicle/DMSO vehicle (*n* = 8), DMSO vehicle/10 mg/kg morphine (*n* = 11), 3 mg/kg IBNtxA/DMSO vehicle (*n* = 11), 3 mg/kg IBNtxA/0 mg/kg morphine (*n* = 11).

##### Testing

In order to reduce novelty-induced locomotor activity, all animals were exposed to the open field twice over two days prior to testing. During these 55-minute pre-exposures, animals received a saline injection after 15 minutes in the open field to familiarize them with the handling and injection process. Behavior during these pre-exposures was not recorded.

On the test day, animals were given an i.p. pre-treatment injection (3 mg/kg IBNtxA or DMSO vehicle) and placed in the open field for 15 minutes. Behavior during this period was not recorded. After 15 minutes, animals were given an i.p. post-treatment injection (10 mg/kg morphine or DMSO vehicle). Locomotor activity was recorded for 40 minutes.

##### Analysis

Locomotor activity for each animal was recorded in 5-minute bins. Overall locomotor activity is represented as the total distance traveled following the second injection.

#### Naloxone-precipitated morphine withdrawal

##### Justification

To determine whether IBNtxA might alter morphine withdrawal-mediated behavior, we tested whether 3 mg/kg IBNtxA could induce any signs of withdrawal or alter the behavioral responses of naloxone-precipitated withdrawal.

##### Treatment groups

16 drug-naïve male C57BL/6 mice were used for this study. All animals underwent the same chronic morphine dosing, then were randomly assigned to two treatment groups for the final test, each receiving a pre-treatment and a post-treatment injection: DMSO vehicle/30 mg/kg naloxone (*n* = 8), and 3 mg/kg IBNtxA/30 mg/kg naloxone (*n* = 8).

##### Treatment and testing

All mice were treated with nine injections of 50 mg/kg morphine over four days (0900 and 1600 hours each day). On day 5, mice received an additional injection of 50 mg/kg morphine in the morning. To observe whether IBNtxA could induce morphine withdrawal symptoms, 8 animals received 3 mg/kg IBNtxA and 8 animals received vehicle injections 3 hours after the final morphine injection. All animals were observed for signs of morphine withdrawal-induced jumping behavior by two observers for 10 minutes while placed inside a 2 L glass beaker. 15 minutes after the IBNtxA or vehicle treatment, all animals received 30 mg/kg naloxone and were again observed for withdrawal-induced jumping behavior by two observers for 10 minutes. For both observation periods, a jump was defined as all four paws leaving the bottom of the beaker. Other signs of opioid withdrawal (e.g., shakes, diarrhea) were looked for but not observed to any substantial degree and are thus not reported. Between every animal, and before and after all training and testing procedures, each glass beaker was cleaned with 70% isopropyl alcohol and allowed to dry completely.

#### Statistical analyses

All data are presented as means ± SEM and were analyzed in GraphPad Prism 6 (San Diego, CA, USA). For analysis of variance (ANOVA) tests, whenever a significant main effect was found, individual group comparisons were carried out using pre-planned Bonferroni t tests.

## RESULTS

### IBNtxA is antinociceptive

In the hot plate assay, 3 mg/kg IBNtxA, but not 1 mg/kg IBNtxA, significantly increased the time to nocifensive movements (Fig. 2A). 3 mg/kg IBNtxA achieved a peak efficacy (∼61 %MPE) comparable to 10 mg/kg morphine (∼64 %MPE), but showed a shorter duration of effect. Individual one-way repeated-measures ANOVAs comparing post-injection treatments against an average of the two baseline (BL) measures revealed significant treatment effects of 10 mg/kg morphine (F(6, 30) = 3.436, p = 0.0106) and 3 mg/kg IBNtxA (F(4, 20) = 5.780, p = 0.0029) treatments, but not 1 mg/kg IBNtxA (F(4, 20) = 0.4840, p = 0.7473). Pre-planned Bonferroni tests determined that 3 mg/kg IBNtxA significantly differed from baseline at 30 minutes post-injection (t > 3.8 and p < 0.01), and 10 mg/kg morphine significantly differed from baseline at 30 and 45 minutes post-injection (t > 3.2 and p < 0.05 for both).

**Figure 1.**
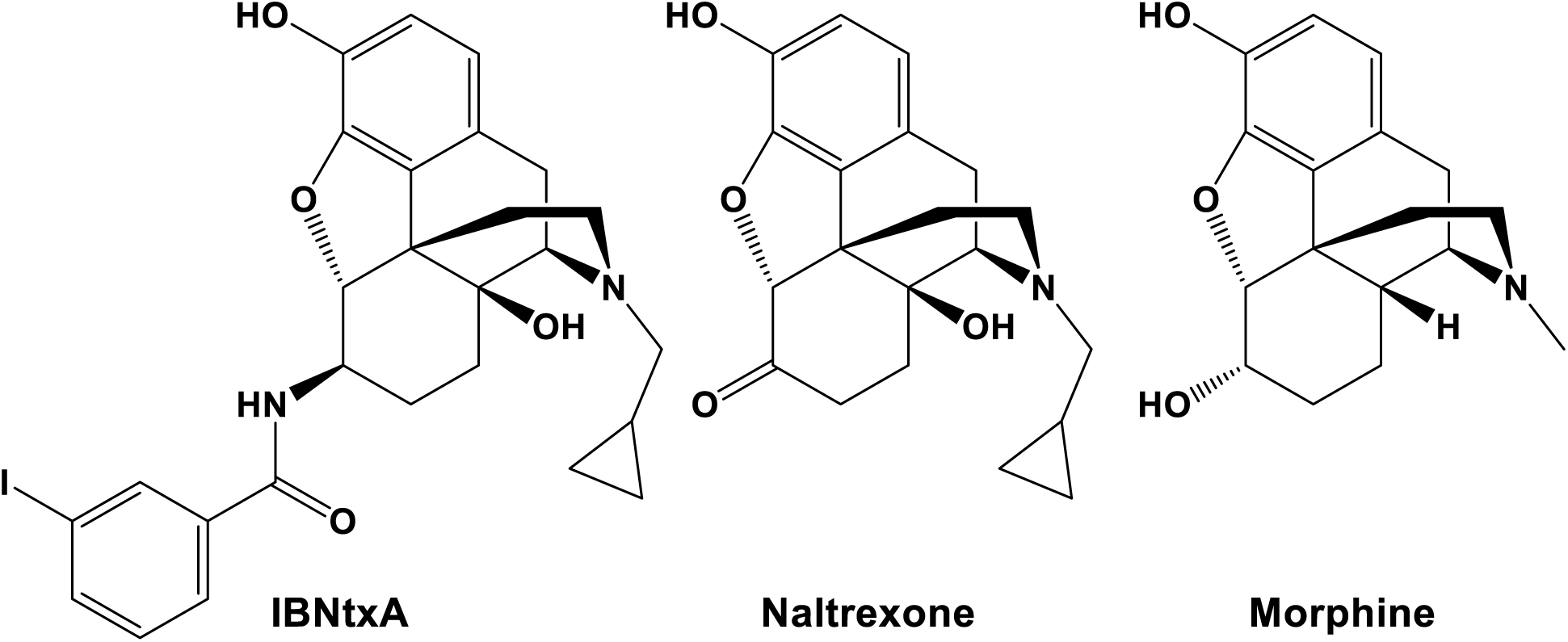
Structures of IBNtxA, naltrexone, and morphine.

**Figure 2.**
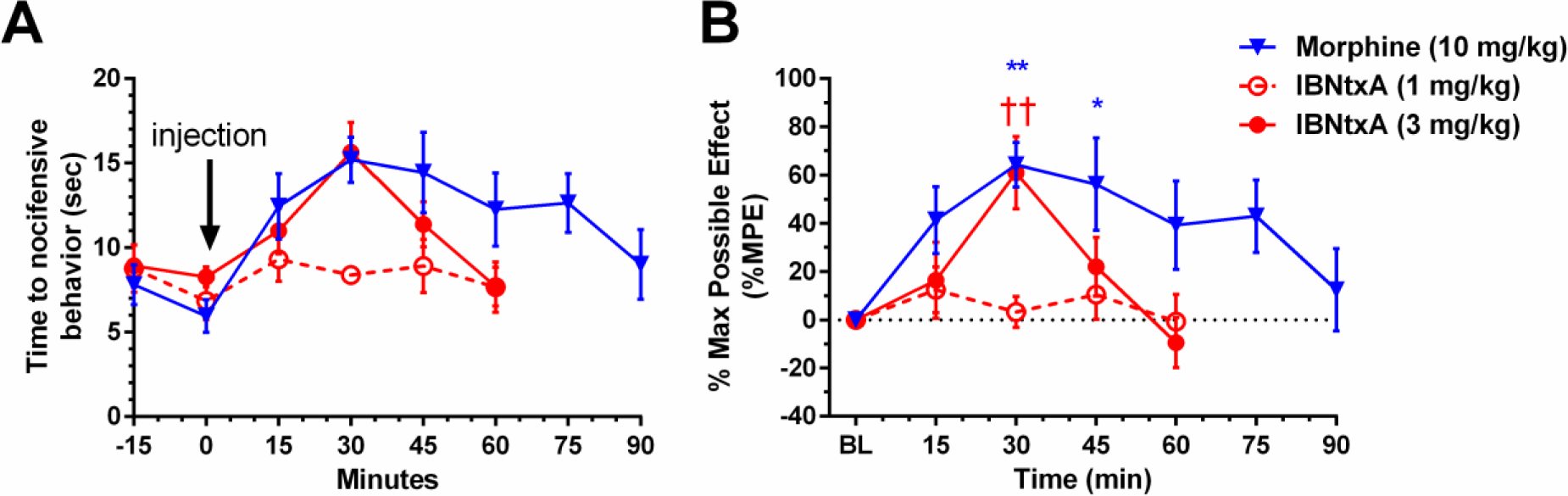
Time-course of hot plate analgesia tests. Male C57BL/6 mice (*n* = 6/group) received IBNtxA or morphine at the indicated doses using a hot plate apparatus set at 55 °C. A. The time elapsing to the first nocifensive response (jumping, paw licking or fluttering) was scored. A maximal latency of 20 s was used to minimize any tissue damage. B. Results analyzed as percent maximal effect (%MPE) for each mouse. All results are presented as means ± SEM; * p < 0.05, ** p < 0.01 compared to BL of morphine; †† p < 0.01 compared to BL of IBNtxA.

### Drug discrimination reveals a complex opioid receptor pharmacology for IBNtxA

Male C57BL/6 mice (*n* = 7) were trained to discriminate 3 mg/kg IBNtxA from DMSO vehicle. IBNtxA proved to be a weak discriminative stimulus, taking nearly 90 training days for all animals to meet training criteria (> 80% correct responses). Initial experiments revealed that drug trials in which animals could respond for 30 minutes substantially disrupted subsequent discrimination training and required extended re-training between drug tests. Therefore, drug tests were redesigned to count only the initial (unrewarded) nose poke after test drug delivery 15 minutes prior to the start of the trial. Only results from this second iteration are presented in Figure 3; panel A indicated the first nose poke response for each tested drug dose and panel B indicates the time to initial nose poke response, an indication of whether a drug dose was behaviorally disruptive. Initial test trials with DMSO vehicle and 3 mg/kg IBNtxA (black squares) indicated that this method could reliably distinguish vehicle from training drug. Dose responses were then recorded for IBNtxA (0.3-3 mg/kg), MOR agonist morphine (0.33-10 mg/kg), KOR agonist U-50488 (0.33-10 mg/kg), DOR agonist SNC162 (3-18 mg/kg), NOP agonist SCH 221510 (1-10 mg/kg), and MOR partial agonist/NOP agonist buprenorphine (0.10-1 mg/kg). To test generalizability, the non-opioid cocaine (3-10 mg/kg) was tested as well.

**Figure 3.**
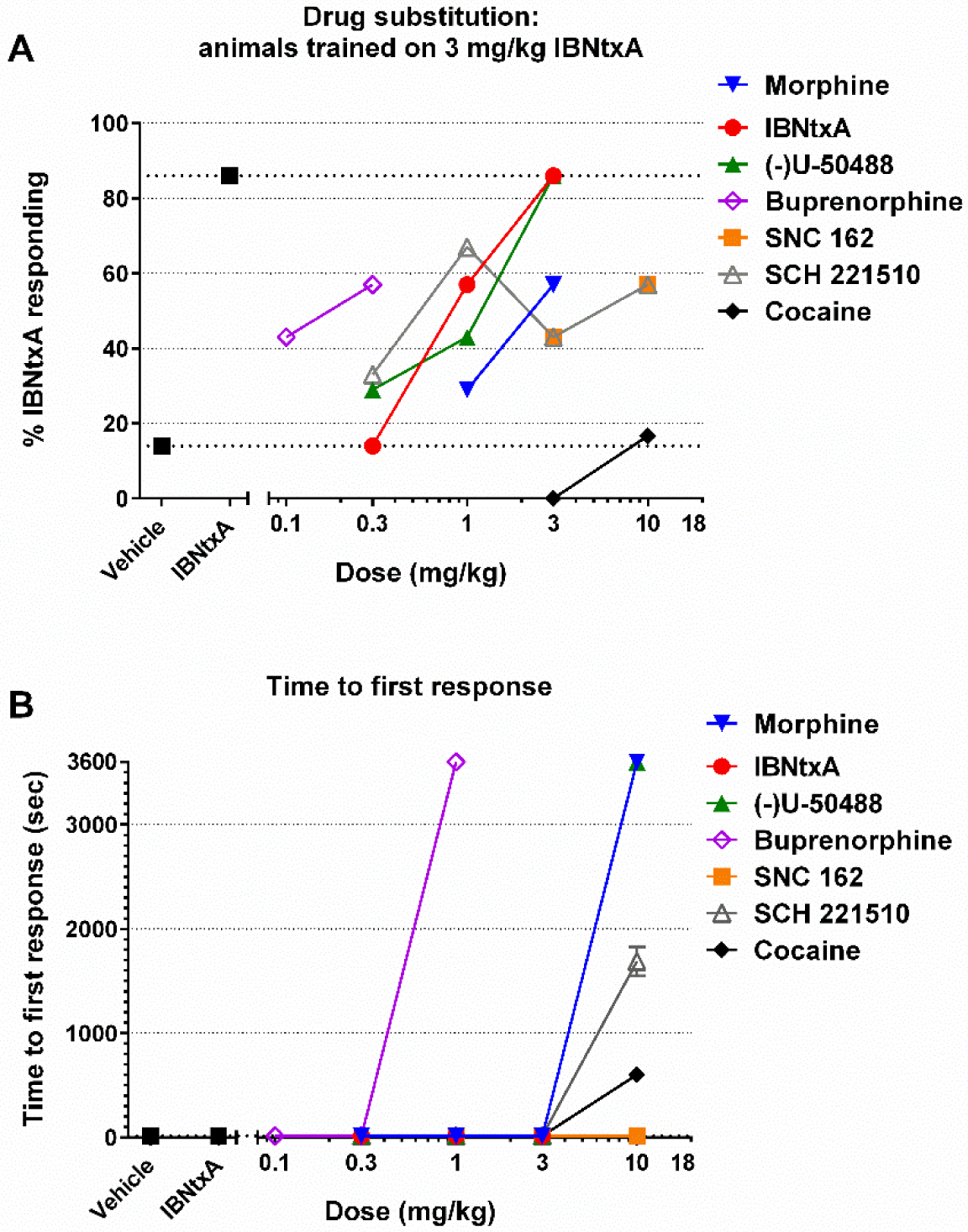
Drug discrimination results for animals trained to distinguish 3 mg/kg IBNtxA from vehicle. A. IBNtxA and KOR agonist U-50488 fully substitute for IBNtxA. MOR agonist morphine, DOR agonist SNC162, NOP agonist SCH 221510, MOR partial agonist/NOP agonist buprenorphine partially substitute for IBNtxA. The psychostimulant cocaine does not substitute for IBNtxA. Because these data ultimately represent the proportion of mice who chose the IBNtxA-paired nose poke hole for their first response, there are no error bars. B. Behavioral disruption of drug responding as determined by the time to first drug response. The highest tested doses of buprenorphine, morphine, SCH 221510, and cocaine each induced a substantial behavioral disruption. These results are presented as means ± SEM.

IBNtxA dose-dependently and fully substituted for itself. MOR agonist morphine, DOR agonist SNC162, NOP agonist SCH 221510 and MOR partial agonist/NOP agonist buprenorphine each partially substituted for IBNtxA, while the psychostimulant cocaine did not substitute for IBNtxA (Figure 3A). At the highest doses tested, morphine (10 mg/kg), SCH 221510 (10 mg/kg), and buprenorphine (1 mg/kg) each disrupted responding (Figure 3B). KOR agonist U-50488 fully substituted for IBNtxA, indicating that KOR signaling effects are likely crucial to the *in vivo* characteristics of IBNtxA.

### IBNtxA does not induce a place preference

To determine whether IBNtxA may have abuse liability, it was tested in an unbiased CPP assay. Seven groups of C57/Bl6 mice (*n* = 12-22 per group) were tested with different doses of IBNtxA, morphine, or DMSO vehicle as the training drug in a three-compartment apparatus. The overall place preference for each drug dose was determined using a preference score (ΔΔsec). A one-way ANOVA comparing all results indicated a significant difference between treatment groups (F(6, 101) = 2.444; p = 0.0301). Pre-planned Bonferroni tests determined that 10 mg/kg morphine significantly differed from vehicle (t > 3.8 and p < 0.05) but no other doses were significantly different from vehicle. In an alternate analysis, using a one-sample t-test to evaluate whether each treatment was significantly different from a hypothetical 0 value that would result from no compartment preference, 10 mg/kg morphine (t(21) = 3.737, p = 0.0012) and 20 mg/kg morphine (t(11) = 3.841, p = 0.0027) significantly differed from 0; all other doses were not significantly different from 0.

### IBNtxA pre-treatment disrupts acquisition of morphine place preference

To further probe the effects of IBNtxA, we investigated whether IBNtxA could disrupt morphine-mediated behaviors. In this experiment, three groups (*n* = 10 each) of C57/Bl6 mice underwent CPP training for 10 mg/kg morphine as above, however 15 minutes prior to each morphine training mice were pre-treated with IBNtxA (1 mg/kg or 3 mg/kg) or vehicle, and all groups received vehicle pretreatment prior to vehicle side training. IBNtxA dose-dependently disrupted acquisition of morphine place preference (Figure 5). One-way ANOVA revealed a significant overall effect of pre-treatment (F(2, 27) = 3.640, p = 0.0399). Pre-planned Bonferroni tests determined that 3 mg/kg IBNtxA significantly differed from vehicle (t > 2.55 and p < 0.05).

**Figure 4.**
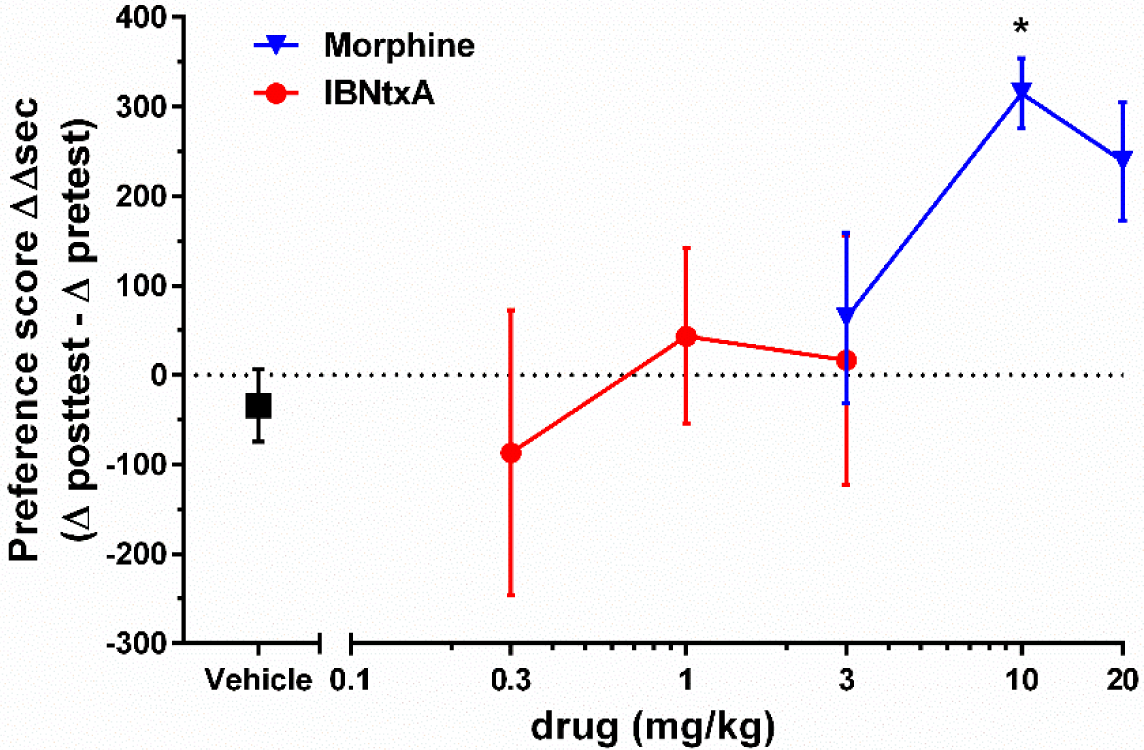
Conditioned place preferences of varying doses of morphine or IBNtxA compared to vehicle-only treatment. 10 mg/kg morphine significantly induced a preference for the drug-paired compartment while IBNtxA did not produce a significant place preference at doses up to 3 mg/kg. All results are presented as means ± SEM; * p < 0.05 compared to vehicle.

### IBNtxA attenuates acute morphine-induced hyperlocomotion

To further evaluate behavioral IBNtxA disruption of morphine-mediated behaviors was behaviorally disruptive, we measured the locomotor activity of mice pretreated with either 3 mg/kg IBNtxA or vehicle followed by either 10 mg/kg morphine or vehicle (Figure 6). Two-way repeated-measures ANOVA revealed significant effects of drug treatment (F(3, 37) = 20.92, p < 0.0001) and time (F(7, 259) = 9.375, p < 0.0001), with a significant interaction (F(21, 259) = 15.05, p < 0.0001). Morphine induced a substantial hyperlocomotion (Figure 6A); pre-planned Bonferroni tests determined that the vehicle/morphine treatment group significantly differed from all other groups at the 15-40 min time points (t > 4.976 and p < 0.0001 for each point) with no other significant differences between points. Two-way ANOVA analysis of total locomotor activity over the 40-minute trial (Figure 6B) revealed significant effects of IBNtxA vs. vehicle pre-treatment (F(1, 37) = 29.19, p < 0.0001) and morphine vs. vehicle post-treatment (F(1, 37) = 7.001, p = 0.0119), with a significant interaction (F(1, 37) = 22.02, p < 0.0001). Pre-planned Bonferroni tests determined that the vehicle/morphine treatment group significantly differed from all other groups (t > 5.427 and p < 0.0001), but all other comparisons were not significant.

**Figure 5.**
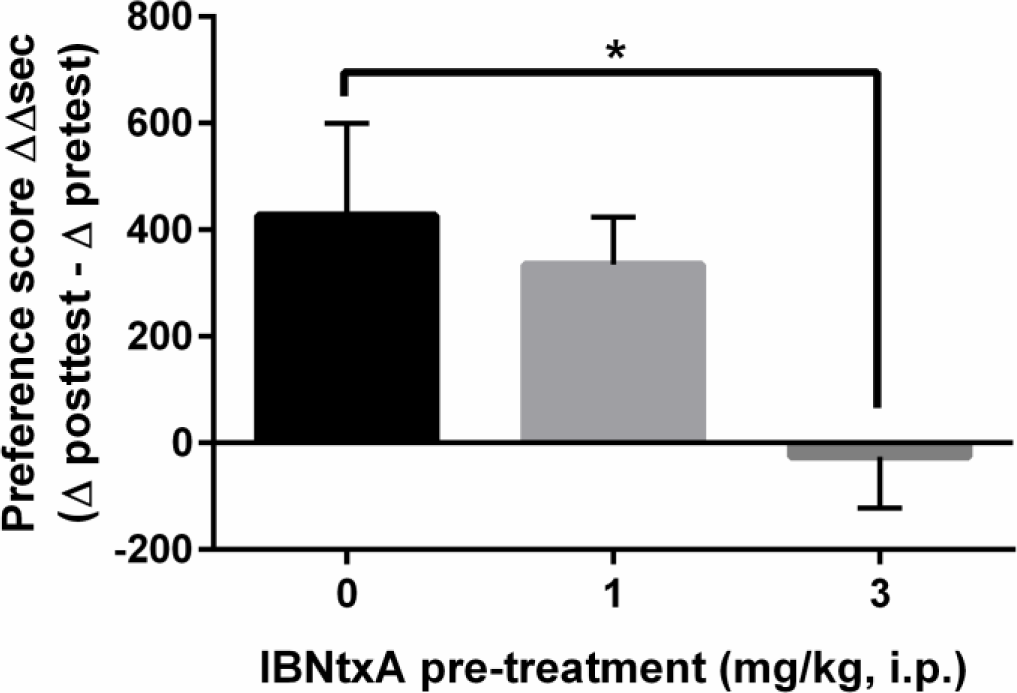
IBNtxA pretreatment attenuated acquisition of morphine CPP. CPP training for 10 mg/kg morphine was modified to include an IBNtxA or vehicle pretreatment prior to standard side training. 3 mg/kg IBNtxA pretreatment significantly attenuated acquisition of morphine-induced place preference. All results are presented as means ± SEM; * p < 0.05 compared to vehicle pre-treatment.

### IBNtxA does not alter naloxone-precipitated morphine withdrawal

To determine whether IBNtxA might alter morphine withdrawal-mediated behavior, we tested whether 3 mg/kg IBNtxA could induce any signs of withdrawal or alter the behavioral responses of naloxone-precipitated withdrawal. 16 mice were treated with nine injections of 50 mg/kg morphine over five days (0900 and 1600 hours except for day 5). 3 hours after the final morphine injection, 8 animals received 3 mg/kg IBNtxA and 8 animals received vehicle injections, and all animals were observed for jumping and other signs of morphine withdrawal for 10 minutes (Figure 7, left side). No withdrawal signs were seen in either group, indicating that 3 mg/kg IBNtxA did not precipitate withdrawal. 15 minutes after the vehicle or IBNtxA pretreatment, all animals received 30 mg/kg naloxone and were again observed for jumping and other signs of morphine withdrawal for 10 minutes (Figure 7, right side). Two-way repeated-measures ANOVA revealed a significant effects of naloxone treatment on jumping behavior (F(1, 14) = 34.62, p < 0.0001), but not IBNtxA treatment (F(1, 14) = 0.2844, p = 0.6022), with no significant interaction (F(1, 14) = 0.2844, p = 0.6022). Pre-planned Bonferroni tests determined that there were no significant differences between vehicle- and IBNtxA-treated groups before or after naloxone treatment (t < 0.76 and p > 0.05 for all comparisons).

**Figure 6.**
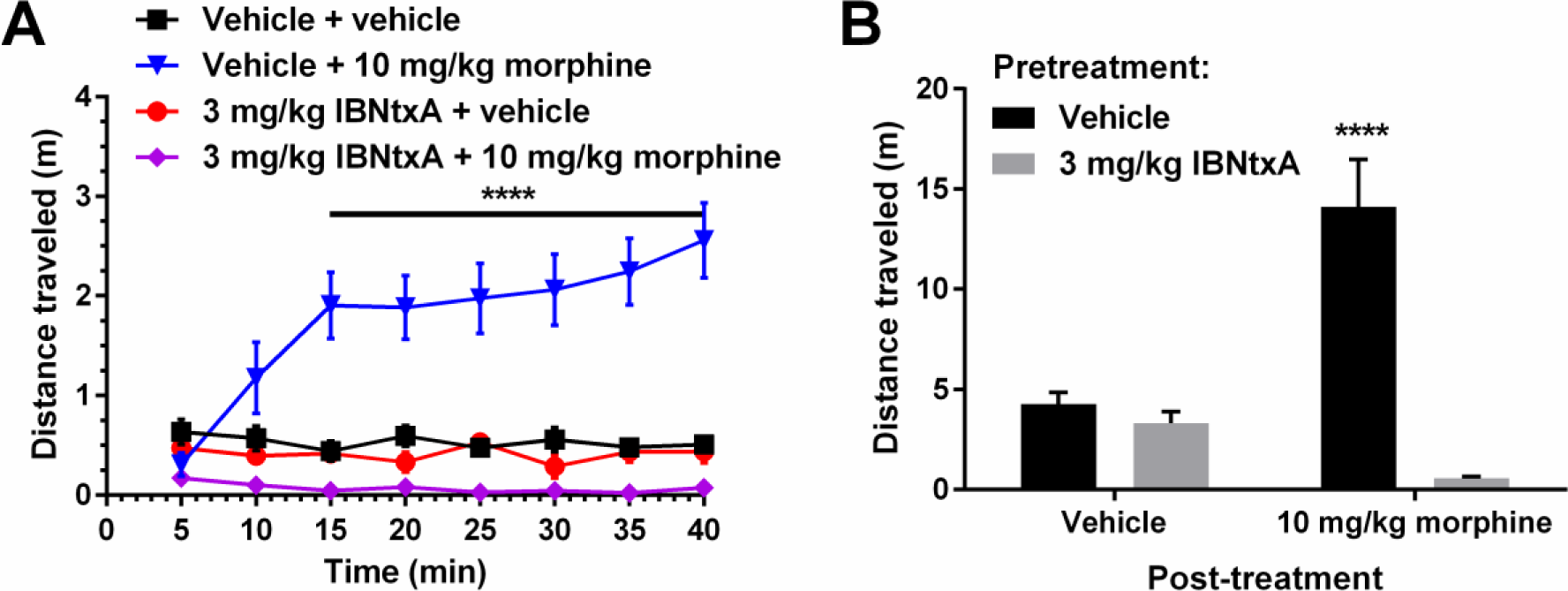
Male mice (*n* = 8-11/group) received DMSO vehicle or 3 mg/kg IBNtxA pretreatments 15 minutes prior to DMSO vehicle or 10 mg/kg morphine administration. A. Timecourse of locomotor activity immediately following the second injection. B. Overall distance traveled of each group. IBNtxA pretreatment significantly attenuated morphine-induced hyperlocomotion but did not significantly alter locomotion following vehicle injection. All results are presented as means ± SEM; **** p < 0.0001 compared to all other groups.

**Figure 7.**
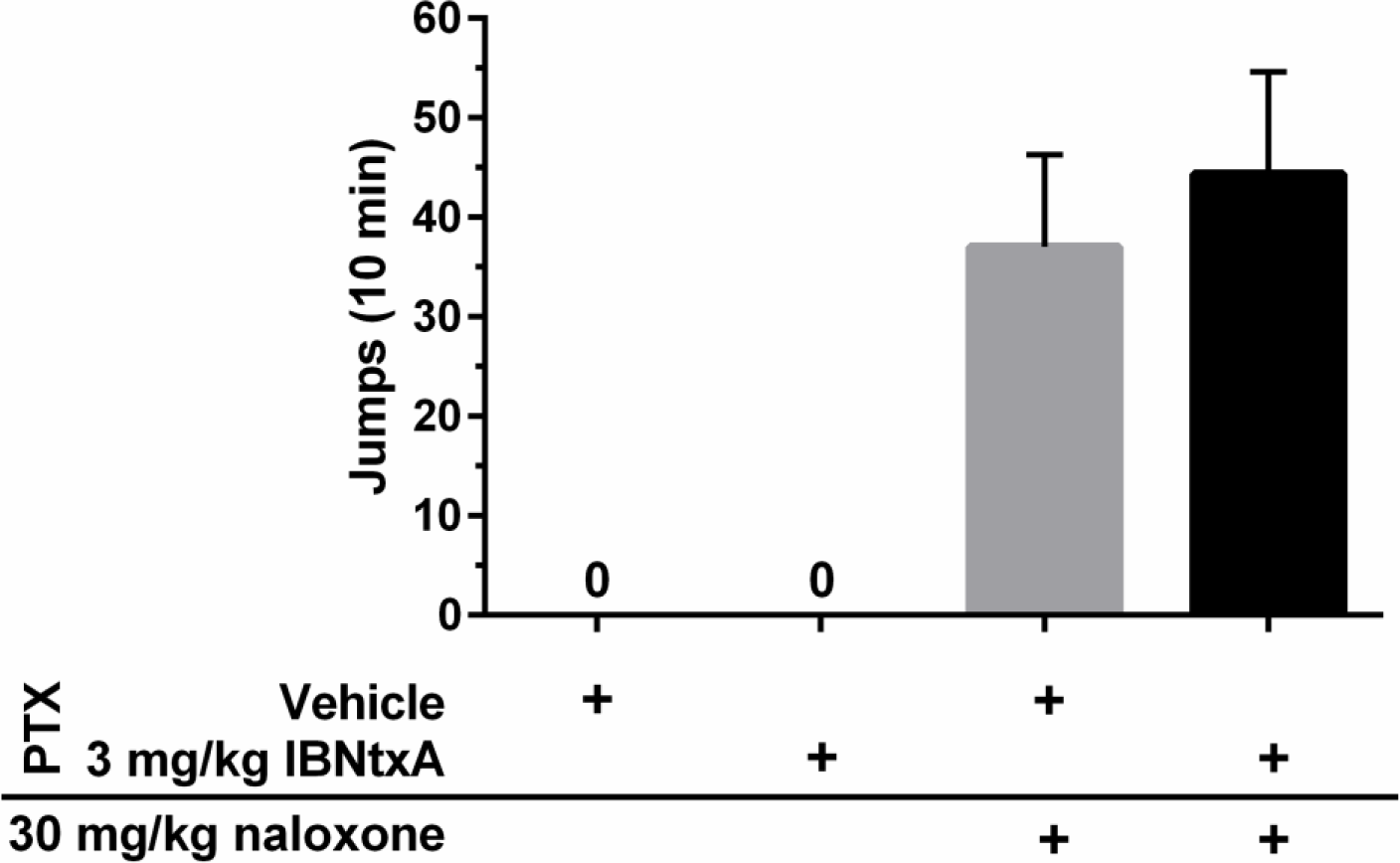
Male mice (*n* = 8/group) received 50 mg/kg morphine. To test whether IBNtxA might induce or induce withdrawal, or blunt naloxone-induced withdrawal, one group received a DMSO vehicle or 3 mg/kg IBNtxA pretreatment. 15 minutes later, both groups received 50 mg/kg naloxone. Jumps were counted over 10 min, defined as all four paws leaving the floor of a glass cylinder. IBNtxA administration did not induce withdrawal or attenuate naloxone-induced withdrawal jumping behaviors.

## Discussion

Since the cloning of the MOR gene, *Oprm1*, multiple splice variants have been discovered. (Pasternak 2004) The functional consequences of MOR variants are poorly understood, but certain splice variants have been reported to differ in subtype localization within the central nervous system and in pharmacological properties, including variable affinity for MOR ligands (Abbadie et al. 2000; Choi et al. 2006; Oldfield et al. 2008; Xu et al. 2014b). The expression of these variants can also be differentially regulated by sex and drug exposure history (Pan et al. 2009; Verzillo et al. 2014). Morphine tolerance is also associated with differential regulation of multiple MOR splice variants (Xu et al. 2015). Because there are no ligands currently available with greater specificity for 6TM/E11 MOR splice variants, IBNtxA is the best current tool for probing the behavioral effects of 6TM/E11 agonism.

Previous IBNtxA studies suggested an intriguing avenue for analgesic medications development: specifically targeting 6TM/E11 splice variants. Additionally, these studies provided evidence that 6TM/E11 MOR variants contribute to the analgesic effects for only certain MOR agonists, such as fentanyl, buprenorphine and heroin, but not for morphine (Grinnell et al. 2016; Lu et al. 2015; Majumdar et al. 2011b; Pan et al. 2009). It is not currently known to what degree signaling by 6TM/E11 splice variants mediates drug-seeking and -taking behaviors for opioid drugs.

IBNtxA was readily synthesized from commercially available naltrexone. This synthetic process resulted in a racemic mixture that underwent enantiomeric purification. Only the (-)-isomer of IBNtxA was tested in behavioral studies as this was the enantiomer tested in previous studies (Grinnell et al. 2014; Majumdar et al. 2011a; Majumdar et al. 2011b). To our knowledge, the (+)-isomer of IBNtxA has not been tested for relevant signaling & behavioral effects; (+)-naloxone and (+)-naltrexone have well-described antagonist effects at the toll-like receptor 4 (TLR4), which may also contribute to the abuse-related effects of cocaine or opioid agonists (Bachtell et al. 2015; Hutchinson et al. 2012; Tanda et al. 2016; Yue et al. 2020), so it would be interesting to test whether the structurally similar (+)-IBNtxA may also have TLR4 effects.

Initial reports determined that IBNtxA was a potent analgesic in the mouse tail-flick and hot plate assays, with peak analgesic effects occurring at 1 mg/kg and 2 mg/kg IBNtxA, respectively (Majumdar et al. 2011b). We independently confirmed the analgesic effect of IBNtxA at 3 mg/kg, but not 1 mg/kg, in a mouse hot plate assay; 3 mg/kg IBNtxA was equipotent to 10 mg/kg morphine, with a similar time to peak effect (30 min) but a shorter duration of effect. Because 3 mg/kg IBNtxA was fully analgesic in the supraspinally mediated hot plate assay, we considered this dose in further testing to be centrally active. Partial substitution of IBNtxA by DOR, MOR, and NOP agonists, and full substitution by a KOR agonist in the DD assays indicate that these receptors may each contribute to IBNtxA-mediated analgesia. Agonism of MOR is well-established to produce analgesia, but DOR and KOR activation also produces antinociceptive effects and have been targets for analgesic drug development (Gaveriaux-Ruff and Kieffer 2011; Nagase and Saitoh 2020) (Abdallah and Gendron 2018). NOP ligands have complicated effects on analgesia, depending on route of administration, and can modulate signaling by other opioid receptors (Cremeans et al. 2012; Toll et al. 2016; Varty et al. 2008).

Prior to this study, the discriminative stimulus properties IBNtxA have never been tested. Furthermore, the discriminative stimulus properties of 6TM/E11 agonism are currently unknown, and previous work identified several opioid agonists that may have important effects mediated by 6TM/E11 activation (Majumdar et al. 2011b; Marrone et al. 2016). In order to determine whether IBNtxA could be used to evaluate the subjective effects of 6TM/E11 agonism, its suite of subjective effects must be measured against classical MOR, KOR, NOP, and DOR agonists. The comparison of IBNtxA against buprenorphine—a non-specific opioid receptor partial agonist—was driven by the finding that buprenorphine may derive some of its analgesic effects via 6TM/E11 agonism (Grinnell et al. 2016).

Based on the results of our analgesic and drug discrimination tests, we tested the abuse liability of IBNtxA in an unbiased CPP paradigm at a range of doses (0.1-3.0 mg/kg) that extended up to our centrally active dose of 3 mg/kg (sufficient to produce full analgesia in the hot plate assay). IBNtxA was previously evaluated at only a single dose in a CPP model (Majumdar et al. 2011b). Based on the results of this study, IBNtxA does not appear to have substantial abuse liability, producing no place preference in an unbiased CPP assay at doses up to 3 mg/kg. Results from the drug discrimination study suggest that this may be a result of substantial KOR agonist activity. KOR agonism produces dysphoric effects in humans and aversive effects in rodent CPP models (Chefer et al. 2013; Land et al. 2009; Mucha and Herz 1985; Pfeiffer et al. 1986). NOP activation can also disrupt morphine CPP (Zaveri et al. 2018).

Based on the lack of an IBNtxA-mediated place preference, we explored the possibility that IBNtxA could disrupt morphine-behaviors. Pretreatment with 3 mg/kg but not 1 mg/kg IBNtxA attenuated the acquisition of place preference to 10 mg/kg morphine. This could be due to several factors, including the MOR partial agonist profile of IBNtxA possibly blocking the full agonist effects of morphine. However, because IBNtxA neither induced morphine withdrawal nor attenuated naloxone-precipitated withdrawal, it’s possible that MOR signaling plays only a modest role in IBNtxA-mediated effects. A more likely explanation, based on the drug discrimination study findings, is that IBNtxA’s KOR agonism effects are disrupting morphine behaviors. KOR agonists produce a dysphoric effects that counter MOR agonist-induced place preference (Karkhanis et al. 2017) and the KOR agonist U-50,488 has been previously reported to block morphine place preference (Funada et al. 1993). 3 mg/kg IBNtxA also reversed morphine-induced hyperlocomotion but did not produce a significant decrease in locomotor activity on its own.

Understanding the suite of behavioral effects seen in this study and in previous studies requires consideration of IBNtxA’s lack of subtype selectivity. IBNtxA has nanomolar affinity *in vitro* for 7TM MOR, 6TM/E11 MOR, KOR, and DOR (Majumdar et al. 2011b). In a prior study, IBNtxA analgesia was partially blocked by nonselective opioid antagonist naloxone, KOR antagonist norbinaltorphimine, and DOR antagonist naltrindole, and fully blocked by MOR antagonist/KOR agonist levallorphan, suggesting potential roles for MOR, KOR, and DOR activation mediating IBNtxA-induced analgesia (Majumdar et al. 2011b). Follow-up studies using genetically modified mice indicated that *in vivo* analgesia and behavioral responses induced by IBNtxA could be largely attributed to 6TM/E11 MOR splice variant signaling (Lu et al. 2015; Wieskopf et al. 2014). In MOR/KOR/DOR triple knockout animals, IBNtxA-mediated analgesia, but not morphine-mediated analgesia, could be rescued via intrathecal administration of lentivirus containing 6TM/E11 MOR splice variants (Lu et al. 2015; Lu et al. 2018). While these results support the hypothesis that 6TM/E11 MOR splice variants have distinct receptor signaling properties and agonist binding capabilities—thus making them potential druggable targets—the prevalence of *substantial* 6TM/E11 expression in wild-type animals is of some dispute. For example, Xu et al. (2014a) reported variable expression of 6TM/E11 splice variants across brain regions and mouse strains using qPCR, but Iadarola et al. (2015) found little evidence of exon 11-containing splice variants in mouse dorsal root ganglion cells using RNA-Seq. Future studies using MOR exon 11 knockout mice (Pan et al. 2009) could more fully dissect the role of 6TM/E11 MOR splice variants in mediating the discriminative properties of IBNtxA and other opioid drugs.

Several limitations are worth noting in this study. First, all tests reported herein were performed on only male mice. There is an established literature that shows important sex-mediated differences in opioid receptor expression and signaling, manifesting in differential effects in tests of analgesia in humans and rodents and important differences in rodent models of drug abuse and relapse (Becker and Chartoff 2019; Craft 2008; Dahan et al. 2008; Lee and Ho 2013). Given the apparent full KOR agonism of IBNtxA in vivo, there are important sex differences in KOR function that may impact analgesia and abuse liability (Chartoff and Mavrikaki 2015). A recent study reported differences between adult male and female C57BL/6 mice in mRNA expression patterns of 6TM/E11 MOR splice variants (Liu et al. 2018); if 6TM/E11 MOR signaling plays an important role in IBNtxA-mediated behavioral effects in wild-type mice—a hypothesis that could not be directly tested in these studies—male and female mice may have differing sensitivity to IBNtxA. Finally, all experiments were performed in the light part of the light/dark cycle, which is the inactive part of the mouse diurnal cycle. Opioid receptor expression levels can vary through the diurnal cycle (Mitchell et al. 1998) and opioid receptor activation can itself alter circadian rhythms (Pacesova et al. 2015; Webb et al. 2015). Thus, it is possible different behavioral effects would be observed if these tests were repeated in the dark cycle.

In conclusion, these studies demonstrate that IBNtxA is a potent analgesic, with a complicated *in vivo* opioid receptor pharmacology in wild-type mice. IBNtxA appears to lack abuse liability at doses up to 3 mg/kg, possibly due to its prominent KOR signaling effects, a possible mechanism by which IBNtxA may disrupt morphine-mediated behaviors.

## Supporting information

Supplementary information - IBNtxA synthesis

## ACKNOWLEDGEMENTS

This work was supported by 1R03DA041560 and Rowan University. The authors thank Ella Babenko, Gun Choi, Konstantinos Totolos, Donald McKay, Falisha Lormejuste, Dylan Souder, Maria Oliveira, Nyesha White, Taylor LaCorte, Peace Nwankwo, Charmayne Lash, Evan Winslow, Andrew Kalash, and Ian Hansen for technical contributions to these studies.

Thanks to Gavril Pasternak and Susruta Majumdar for initial guidance on these studies and for providing samples of IBNtxA used in pilot studies and in validation of in-house IBNtxA synthesis.

