## Supplementary information - IBNtxA synthesis for "Abuse liability, antinociceptive, and discriminative stimulus properties of IBNtxA"

**SUPPLEMENTARY MATERIAL**

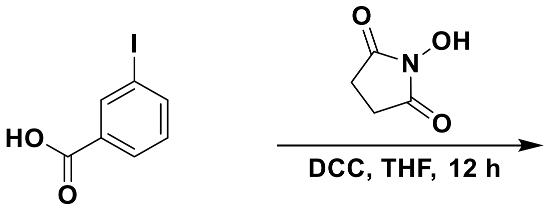


**2,5-Dioxopyrrolidin-1-yl 3-iodobenzoate (S1):** 3-Iodobenzoic acid (2.00 g, 7.90 mmol, 1.0 equiv) was dissolved in anhydrous THF (18 mL, 0.45 M) and cooled to 0 °C on an ice-bath before addition of DCC (1.81 g, 8.69 mmol, 1.1 equiv) as a solid in one portion (a white precipitate formed rapidly). N-hydroxylsuccinimide (1.00 g, 8.69 mmol, 1.1 equiv) was then added a solid in one portion. The cooling bath was removed and the reaction stirred at ambient temperature for 12 hours. At this point, the suspension was filtered and the solids were washed with THF. The filtrate was concentrated to afford a white solid which was purified by chromatography on silica gel (1:9 to 1:4; EtOAc-heptanes) to afford the activated ester **S1** (2.23 g, 82%) as a white solid. $^{1}$**H-NMR** (400 MHz, $CDCl_3$) δ 8.44 (d, *J* = 1.9 Hz, 1H), 8.09 (dd, *J* = 8.0, 1.5 Hz, 1H), 8.00 (dd, *J* = 7.9, 1.6 Hz, 1H), 7.27 (t, *J* = 7.9 Hz, 1H), 3.05 – 2.77 (m, 4H); $^{13}$**C-NMR** (100 MHz, $CDCl_3$) δ 169.0, 160.4, 143.6, 138.9, 130.4, 129.4, 126.8, 93.9, 25.5.

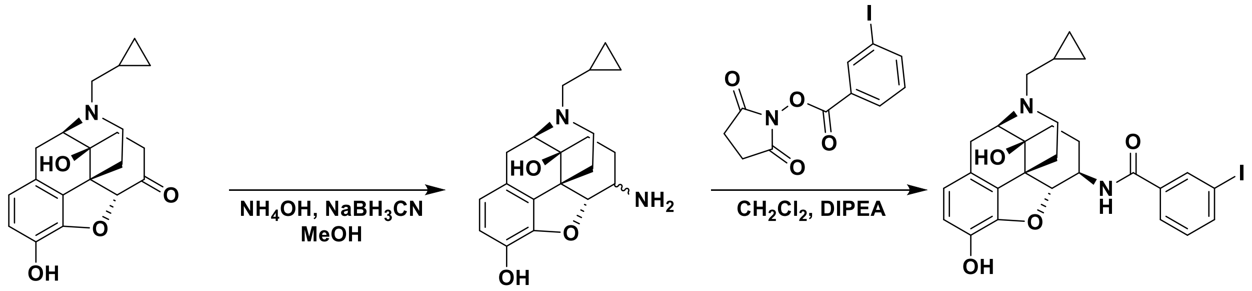


***N*-((4R,4aS,7R,7aR,12bS)-3-(cyclopropylmethyl)-4a,9-dihydroxy-2,3,4,4a,5,6,7,7a-octahydro-1H-4,12-methanobenzofuro[3,2-e]isoquinolin-7-yl)-3-iodobenzamide (IBNtxA):** A vial was charged with naltrexone hydrochloride (1.0 equiv) and ammonium acetate (10 equiv) in MeOH (0.6 M) and the mixture was stirred for 10 minutes before addition of sodium cyanoborohydride (0.7 equiv) as a solution in MeOH. The mixture was stirred for 12 hours before being quenched with 1 M NaOH (0.3 mL). The volatiles were removed *via* rotary evaporation to provide a residue that was diluted with water (2 mL), and extracted with $CH_2Cl_2$. Drying over $Na_2SO_4$ and concentration afforded the crude naltrexamine which was then dissolved in anhydrous $CH_2Cl_2$ (0.1 M) and treated with the NHS-activated ester **S1** (1.1 equiv) and DIPEA (1.1 equiv). This mixture was stirred for 2 hours at ambient temperature before being diluted with $CH_2Cl_2$ and washed with water to give a residue which was purified by chromatography on silica gel (1:99 to 1:20; MeOH-$CHCl_3$) which gave a small forerun of the *N,O*-bisacylated product, followed by the α-isomer, and finally by **IBNtxA** (typically isolated in 23-31 % yield) as a white solid. The purified product was converted to its hydrochloride salt by dissolution of the freebase in diethyl ether and addition of 4 M HCl in diethyl ether. The white precipitate was collected by filtration and dried to give **IBNtxA•HCl** as a white solid (68% yield): **MP:** 203−207 °C. **$[\alpha]_D$** = −161.2 (c 0.1, $CHCl_3$). **$^1$H-NMR** (400 MHz, $CDCl_3$) δ 8.17 (s, 1H), 7.9 (d, J = 8.8 Hz, 1H), 7.76 (d, J = 8.8 Hz, 1H), 7.16−7.12 (m, 1H), 6.68 (d, J = 10.6 Hz, 1H), 6.57 (d, J = 10.6 Hz, 1H), 5.92 (m, 1H), 5.25−5.18 (m, 2H), 4.57 (d, J = 8.8 Hz, 1H), 4.13 (m, 1H), 3.14−1.25 (m, 14H). **$^{13}$C-NMR** (100 MHz, $CDCl_3$) δ 165.4, 142.9, 140.3, 139.3, 136.6, 136.2, 135.3, 130.8, 130.2, 126.3, 124.9, 119.3, 118.1, 117.6, 94.3, 92.9, 70.2, 62.4, 57.8, 50.5, 47.3, 43.6, 31.5, 29.0, 23.2, 22.7 ppm. **ESI-MS** m/z (rel int): (pos) 559.1 ($[M+H]^+$, 100). **HRMS** (ESI): Calculated for: $C_{26}H_{28}N_2O_4I^+$: 559.1094; found, 559.1088.
